# Thermodynamic and mechanistic insights into coupled binding and unwinding of collagen by matrix metalloproteinase 1

**DOI:** 10.1101/2020.08.28.272104

**Authors:** Szymon W. Manka, Keith Brew

**Affiliations:** Nuffield Department of Orthopaedics, Rheumatology and Musculoskeletal Sciences, Kennedy Institute of Rheumatology, University of Oxford, Oxford, UK; Department of Biomedical Sciences, Florida Atlantic University, Boca Raton, FL, USA

**Keywords:** isothermal titration calorimetry, heat capacity change, MMP, collagen unwinding, conformational selection, induced fit, coupled equilibria, collagenolysis

## Abstract

Local unwinding of the collagen triple helix is a necessary step for initiating the collagen degradation cascade in extracellular matrices. A few matrix metalloproteinases (MMPs) are known to support this key process, but its energetic aspects remain unknown. Here, we captured the thermodynamics of the triple helix unwinding by monitoring interactions between a collagen peptide and MMP-1(E200A) – an active-site mutant of an archetypal vertebrate collagenase – at increasing temperatures, using isothermal titration calorimetry (ITC). Coupled binding and unwinding manifests as a curved relationship between the total enthalpy change and temperature of the reaction, producing increasingly negative heat capacity change (ΔΔC_p_ ≈ −36.3 kcal/molK^2^). A specially designed solid-phase binding and cleavage assay (SPBCA) reported strain in the catalytically relevant unwound state, suggesting that this state is distinct from the horizon of sampled conformations of the collagenase-susceptible site. MMP-1 appears to blend selected fit with induced fit mechanisms to catalyse collagen unwinding prior to cleavage of individual collagen chains.

**Graphical Abstract:** 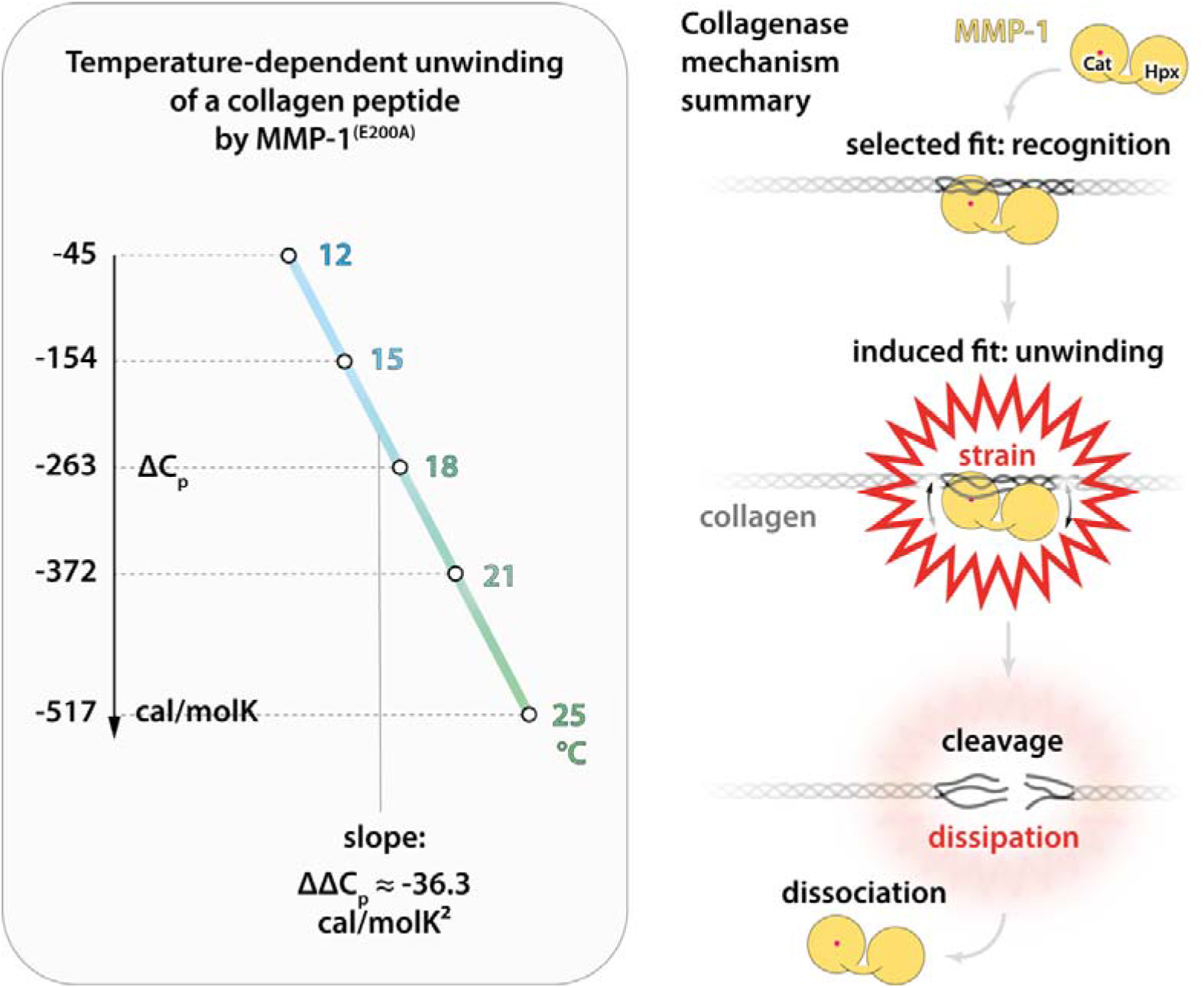

**Highlights:**

- The unwinding of the collagen triple helix by matrix metalloproteinases (MMPs) is critical for collagen catabolism, but how MMPs harness bioavailable energy for this energetically expensive process is enigmatic
- Rising temperature causes linear increases of the negative heat capacity change associated with the interaction of MMP-1 with a collagenase-susceptible triple-helical peptide, indicating enrichment of hydrophobic contacts during the triple helix unwinding
- The complex of MMP-1 with locally unwound collagen holds considerable structural strain, which becomes relieved upon cleavage of the unwound collagen strand(s)
- MMP-1 consumes heat to drive specific (structurally distinct) collagen unwinding by increasing entropy associated with evolving hydrophobic anchor points between the enzyme and the substrate
- The prototypic collagenase MMP-1 appears to blend conformational selection with induced fit, shedding light on the thermodynamic principles by which MMPs trigger collagen breakdown and supporting general mechanistic conclusions concerning conformational changes coupled to protein-protein binding

---

Collagenolysis is an important process in development, organ morphogenesis and tissue repair. When perturbed, it contributes to diseases such as arthritis, atherosclerosis, aneurysm and cancer [1,2]. It is instigated by selected enzymes from the matrix metalloproteinase family (MMP −1, −2, −8, −13, and −14) that share a unique ability to support local unwinding of interstitial triple-helical collagens (types I-III) [3–5]. All collagenolytic MMPs comprise at least two domains, a catalytic (Cat) domain and a hemopexin (Hpx) domain, connected by a flexible linker. Only a single polypeptide chain can fit in the active site cleft of a Cat domain, thus unwinding of the three collagen strands, which absolutely requires input from an Hpx domain, is necessary for the cleavage to occur [6,3,7,8]. The unwinding takes place at a specific site, approximately ¾ of the way from the N-terminus of a collagen molecule, which results in characteristic ¾ and ¼ length cleavage products. These collagen fragments are unstable at body temperature and, as they start to unfold, they become vulnerable to other, less selective proteases. Thus, the initial cleavage by MMPs is the rate-limiting step of collagen breakdown, and its exact mechanism is of wide interest.

A number of hypotheses have been formulated with regard to the mechanism of collagen unwinding [5,9]. The most recent models, focusing on MMP-1 (the prototypic collagenase), agree on several basic principles, but favour different energetic (thermodynamic) explanations for the process, based on two classic concepts: 1) conformational selection, i.e. passive recognition of a pre-existing unwound state, sampled among other fluctuating conformers (selected fit) [10–13] or 2) induced fit, i.e. induction of the functionally relevant unwound state only by the enzyme (while other loosely folded conformers may still be spontaneously sampled) [3,14,15]. Despite the numerous biochemical [3,12,16–19], structural [7,14,20–23], and computational [10,11,15,24] studies, the thermodynamic properties of collagen binding and unwinding by MMP-1 remain unclear.

We used isothermal titration calorimetry (ITC) to obtain a complete thermodynamic profile of the interaction between MMP-1(E200A) (active site mutant, human sequence) and a collagen peptide, which were previously used for co-crystallisation [7] (Fig. 1a and S1). It immediately became apparent that this interaction is endothermic and driven by entropy (Fig. S1a). The total enthalpy (ΔH) and entropy (ΔS) at each recorded temperature (T) exhibit a typical compensation, resulting in approximately constant free energy (ΔG = ΔH - TΔS ≈ −8.6 kcal/mol) (Fig. S1b). However, the absolute values of ΔH and ΔS change much faster than linearly with increasing temperature (Fig. 1a and Fig. S1b).

**Fig 1.**
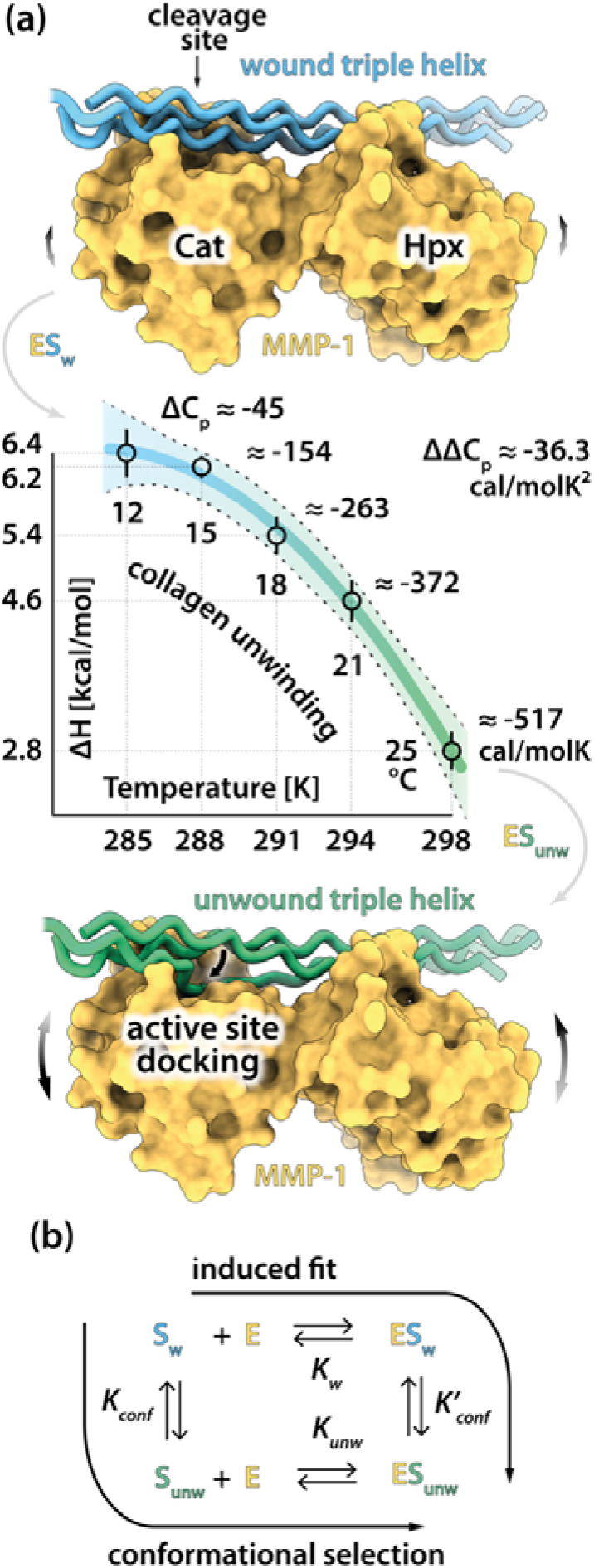
Curved temperature dependence of the observed enthalpy of MMP-1(E200A) interaction with a triple-helical peptide reflects local unwinding of the triple helix. (a) A collagen peptide containing collagenase recognition and cleavage sites, previously co-crystallised with MMP-1(E200A) [7] (top structure: Cat, catalytic domain; Hpx, hemopexin domain), was titrated with MMP-1(E200A) using ITC, as described in Materials and Methods. The apparent change in enthalpy (ΔH) was recorded at increasing temperatures (Fig. S1), revealing a non-linear ΔH vs. T dependence, which has been fitted with eq. 5 (goodness of fit: R^2^ = 0.9970). The fitted curve (solid line) is shown together with its 95% confidence intervals (dashed lines) and is colored with a blue-green gradient to illustrate a changing proportion of wound (blue) vs. unwound (green) states of the collagen peptide bound to the enzyme (ES_w_ vs. ES_unw_) as the temperature increases. The heat capacity change (ΔC_p_) at each temperature is given by eq. 4. It evolves together with the enzyme-substrate interaction, becoming increasingly more negative with increasing temperature, according to ∆∆C_p_ of approx. −36.3 kcal/molK^2^. An illustrative model of the unknown ES_unw_ complex is shown at the bottom. The curved double-headed arrows indicate the inter-domain flexibility of MMP-1, which increases with temperature (arrow size). (b) Coupling between binding and conformational equilibria of the enzyme-substrate complex. S_w_, triple-helical (wound) collagen substrate; S_unw_, unwound collagen substrate; E, enzyme (MMP collagenase); *K*_*w*_ and *K*_*unw*_, intrinsic equilibrium association constants (*K*_*w*_ = [ES_w_]/{[E][S_w_]}, *K*_*unw*_ = [ES_unw_]/{[E][S_unw_]}); *K*_*conf*_ and K′_*conf*_, constants for the conformational equilibrium (*K*_*conf*_ = [S_unw_]/[S_w_] = *K*′_*conf*_{K_w_/K_unw_}; *K*′_*conf*_ = [ES_unw_]/[ES_w_] = *K*_*conf*_{*K*_*unw*_/*K*_*w*_}), square brackets denote molar concentration. Color-coding as in (a).

This strong non-linearity likely corresponds to the well-documented structural change, i.e. the local unwinding of the triple helix accompanying the complex formation [3,14], as illustrated in a simple two-state interaction scheme (Fig. 1b).

Regardless of the pathway in the scheme (induced fit or conformational selection), the data suggest an increasing ratio of the unwound to wound complex ([ES_unw_]/[ES_w_]) with rising temperature (Fig. 1b). Thus, at lower temperatures little or no unwinding occurs (conformational equilibrium constants *K*_*conf*_ or *K*′_*conf*_ ≪ 1 (Fig. 1b)), and the interaction might be described as analogous to lock-and-key, whereas at higher temperatures (up to the melting point of the peptide, T_m_ ≈ 30 °C [7]) the proportion of the unwound complex grows (reaching *K*_*conf*_ or *K*′_*conf*_ ≫ 1), and the thermodynamic contribution of the unwinding becomes increasingly more significant. Consequently, the ΔH vs. T plot is concave (Fig. 1a), because ΔH at any temperature (T) is a combination of the binding term (ΔH_bind_) and the unwinding term (ΔH_unw_) multiplied by the fraction of the unwound complex (f_unw_), which contributed a heat of unwinding [25]:

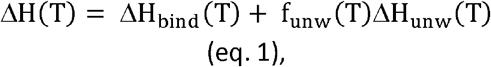

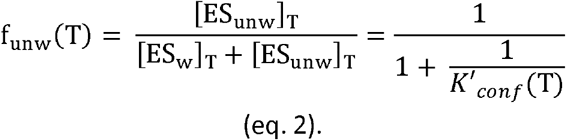

Such coupling of the binding and unwinding equilibria gives rise to temperature-dependence of the apparent heat capacity change (ΔC_p_) of the system [26]:

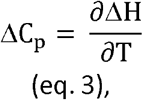

which can be calculated for any temperature (T) using an equation containing a ΔΔC_p_ term that describes a linear change of ΔC_p_ with temperature [27]:

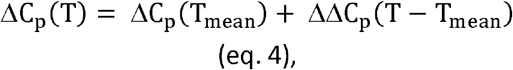

where T_mean_ = 291.2 K (mean temperature within the considered range). The ΔΔC_p_ of approximately −36.3 cal/molK^2^ was obtained by fitting the following function to the ΔH vs. T data (Fig. 1 and S2):

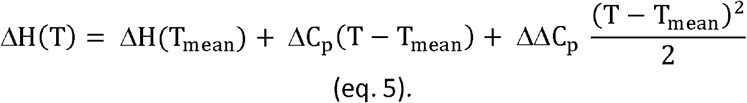

Interestingly, similar ΔΔC_p_ values were reported for DNA binding proteins (e.g. −19 cal/molK^2^ for Klentaq DNA polymerase)[27].

It is broadly accepted that ΔC_p_ is correlated with changes in solvent accessible surface area (ΔASA) upon enzyme-substrate interaction, with burial of polar (ΔASA_pol_) and apolar (ΔASA_apol_) surfaces contributing positively and negatively to ΔC_p_, respectively. Accordingly, the strongly negative ΔΔC_p_ would suggest that the unwinding could be driven by hydrophobic interactions, compensated entropically by the disruption of ordered hydration shells surrounding the relevant hydrophobic patches on both the enzyme and the substrate, prior to interface formation [7]. However, calculation of ΔC_p_ based on the previously proposed parameters [28] and ΔASA_pol/apol_ derived using the GetArea server (http://curie.utmb.edu/getarea.html) from the only available structure of the wound (ES_w_) complex [7] (Fig. 1a, top), gives ΔC_p,calc_ ≈ −387 cal/molK, which correlates best with the experimental ΔC_p_ value observed at 21 °C (ΔC_p_(294 K) ≈ −372 cal/molK, Fig. 1a), where we anticipate prevalence of the unwound (ES_unw_) complex, rather than the wound complex. The latter is likely best represented by the ΔC_p_ value recorded at the lowest temperature (ΔC_p_(285K) ≈ −45 cal/molK, Fig. 1a). Such poor correlations between experimental and ΔASA_pol/apol_-derived ΔC_p_ values are not uncommon for systems exhibiting strongly negative ΔC_p_ and ΔΔC_p_, and may result from incorrect (non-universal) parametrisation [27]. Nonetheless, an overall effect of a conformational change coupled to molecular recognition, such as a qualitative change in ΔASA, may still be inferred from the observed temperature-dependence of ΔC_p_. Here, as previously noted, the particular temperature-dependence of ΔC_p_ hints of a potential enhancement of hydrophobic contacts upon transition from the wound (ES_w_) to the unwound (ES_unw_) complex. This agrees with the previously proposed docking of the P1’ leucine residue of an unwound collagen strand in the hydrophobic S1’ pocket of the enzyme [7] (Fig. 1a, bottom model), prior to peptide bond hydrolysis (P1’ is a residue C-terminally adjacent to a scissile bond and S1’ is its acceptor site in an enzyme [29]).

ITC is useful not only for obtaining thermodynamic signatures, but also for estimating binding affinities (equilibrium association and dissociation constants, *K*_*a*_ and *K*_*d*_ = 1/*K*_*a*_, respectively), from the slope portion of the interaction isotherm (Fig. S1). The affinity of MMP-1(E200A) for its binding site in collagen was previously found to be temperature-dependent over a wide temperature range (4 – 37 °C), employing a solid-phase binding assay (SPBA) using immobilized native collagen [7]. Here, using a collagen peptide, we could only estimate binding affinities for a relatively narrow temperature range (12 – 21 °C) (Fig S2a), because of the thermal instability of the peptide at higher temperatures. We lack data points for the slope of the isotherm already at 25 °C (Fig S1a), where the peptide starts to unfold according to the previously published circular dichroism (CD) data (Fig. S2b, reanalysed data from [7]). The apparent *K*_*d*_ of the MMP-1(E200A)-collagen peptide interaction in our workable temperature range (12 – 21 °C) appears to be constant within the experimental error (0.25 ≤ *K*_*d*_ ≤ 0.5 μM) (Fig. S2a). Since SPBA is a non-equilibrium method, it cannot deliver the *bona fide* equilibrium binding constants for direct comparisons with the ITC-derived values but can be used for relative comparisons. Accordingly, our ITC results are internally consistent with the previously reported SPBA results, showing small differences between the apparent collagen-binding affinities of MMP-1(E200A) at 4 and 20 °C (0.8 ≤ *K*_*d*_ ≤ 1 μM) (Fig. S2c, reanalysed data from [7]). Further, SPBA data suggest that MMP-1(E200A) roughly doubles its affinity for collagen at 37 °C (*K*_*d*_ ≈ 0.4 μM) compared to 20 °C (*K*_*d*_ ≈ 0.8 μM) or 4 °C (*K*_*d*_ ≈ 1 μM) (Fig. S2c). Above 37 °C collagen melts [30] and the affinity of MMP-1(E200A) for it drastically drops [7]. Thus, within the 4 – 37 °C range, the observed *K*_*a*_ of the MMP-1-collagen interaction will be a population-weighted average of the intrinsic association constants of all the states present [26]:

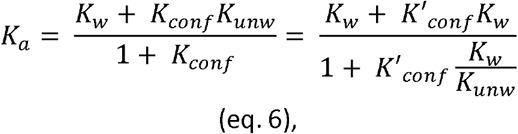

where *K*_*w*_ and *K*_*unw*_ are the intrinsic association constants of ES_w_ and ES_unw_, respectively, and ES_w_ and ES_unw_ are considered boundary states for the conformational continuum (Fig. 1b). Assuming that at 4 °C the ES_w_ population is approximately homogenous (*K*_*conf*_ or *K*′_*conf*_ ≪ 1) and at 37 °C the ES_unw_ population is approximately homogenous (*K*_*conf*_ or *K*′_*conf*_ ≫ 1), then *K*_*a*_(4 °C) ≈ *K*_*w*_ and *K*_*a*_(37 °C) ≈ *K*_*unw*_, hence *K*_*unw*_ ≈ 2*K*_*w*_. Based on this notion, we simulated apparent *K*_*a*_ (and *K*_*d*_) values that would be observed using ITC in our experimental temperature range if *K*_*w*_ and *K*_*unw*_ were fixed at 2 × 10^6^ and 4 × 10^6^ μM, respectively, and *K*_*conf*_ or *K*′_*conf*_ values were changing from 0.1 or 0.2 at 12 °C (5-fold higher [ES_w_] than [ES_unw_]) to 5 or 10 at 21 °C (10-fold higher [ES_unw_] than [ES_w_]), respectively. Such simulated apparent *K*_*d*_ values range from 0.46 μM at 12 °C to 0.27 μM at 21 °C (Fig. S2d), which agrees with our observed values within the experimental error (Fig. S2a). Thus, the intrinsic affinity of MMP-1 for the unwound state of collagen may indeed be 2-fold higher than that for the wound state, even when the observed *K*_*a*_ appears similar over a relatively narrow temperature range, given the uncertainty of our ITC measurements.

Based on the ITC data alone, we cannot discern which model for the dynamic MMP-1-collagen interaction is right: 1) conformational selection or 2) induced fit (Fig. 1b). To gain the relevant insight, we designed a custom solid-phase binding and cleavage assay (SPBCA), in which the binding of MMP-1(E200A) (the *unwinder* enzyme) to immobilized native collagen is monitored in the presence of a *cutter* enzyme, such as an isolated active Cat domain of MMP-1 (MMP-1Cat), supplied *in trans* [3]. Such a *cutter* enzyme can only cleave unwound collagen chains, therefore, its activity (at physiological conditions) requires prior MMP-1(E200A) interaction with the triple-helix [3]. We found that the binding of MMP-1(E200A) to collagen at 20 °C is dose-dependently enhanced in the presence of a potent *cutter* enzyme (MMP-1Cat) [3], as opposed to a poor *cutter* enzyme (MMP-3Cat, the isolated catalytic domain of MMP-3) [31], which showed no effect (Fig. 2a). We then confirmed by Western Blot that collagen is indeed cleaved by MMP-1Cat under the conditions of the assay and that this cleavage stimulates deposition of MMP-1(E200A) on collagen (Fig. 2b and c). These results suggest that the cleavage in the ES_unw_ complex executed by the externally supplied *cutter* activity improves the fit between the *unwinder* enzyme and the clipped substrate, enabling formation of a tighter complex. We wondered if this effect is temperature-dependent, so we repeated the MMP-1Cat treatment for a range of temperatures (4 – 37 °C) (Fig. 2b). As expected, the effect was not observed below 15 °C, where little unwinding occurs [3] (Fig. 1a), and was increasingly more noticeable with increasing temperature, with the peak stimulation observed between 30 °C and 37 °C (Fig. 2d), where the unwinding is optimal. We also checked the time-dependence of this effect at 25 °C, which showed peak stimulation between 2 h and 6 h, and nearly no stimulation after 24 h (Fig. 2e). This suggests that MMP-1(E200A) binding to collagen chains becomes stronger when they are only partially cleaved in the context of the ES_unw_ complex, and not excessively cleaved. For a final negative control, we used collagen in which the collagenase cleavage site was mutated, rendering it uncleavable by MMPs [32]. The binding of MMP-1(E200A) to the non-cleavable collagen was insensitive to the presence of MMP-1Cat at the optimal (30 °C) temperature (Fig. 2f), which ultimately confirmed that the observed binding enhancement of MMP-1(E200A) to collagen depends on the cleavage in the ES_unw_ complex.

**Fig 2.**
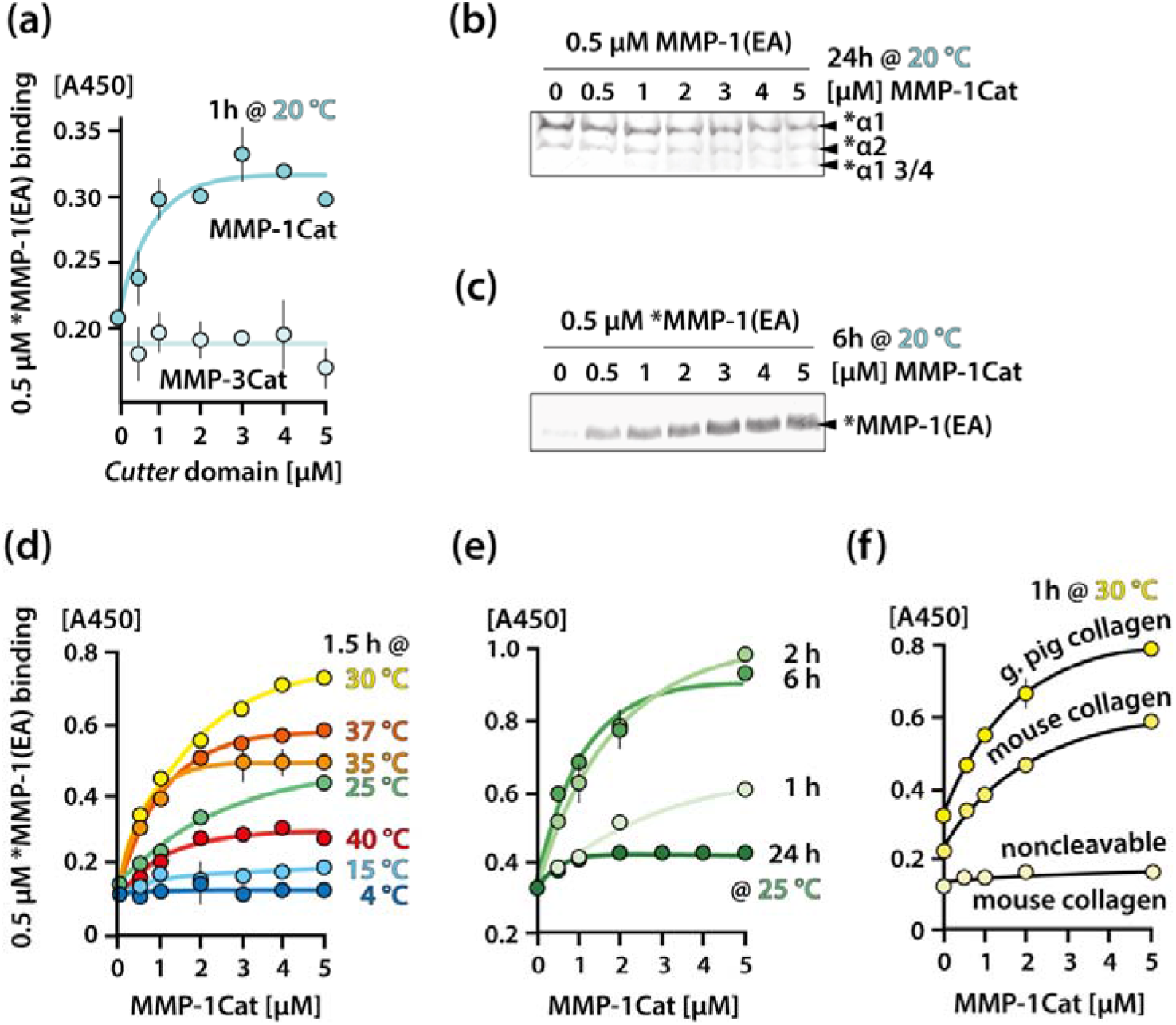
Partial cleavage in the unwound (ES_unw_) complex enhances MMP-1(E200A) interaction with the clipped collagen. (a) Solid-phase binding and cleavage assay (SPBCA) reporting the binding of biotinylated (*) MMP-1(E200A) to collagen in the presence of increasing concentrations of the isolated catalytic domains of MMP-1 (MMP-1Cat) and MMP-3 (MMP-3Cat) (*cutter* domains), over 1 h at 20 °C. (b) Western blot reporting the cleavage of biotinylated (*) immobilized collagen in the presence of MMP-1(E200A) and increasing concentrations of MMP-1Cat, over 24 h at 20 °C. The ¾ collagenase cleavage product of the collagen α1 chain was detected. (c) Western blot reporting the deposition of biotinylated (*) MMP-1(E200A) on immobilized collagen in the presence of increasing concentrations of MMP-1Cat, over 6 h at 20 °C. (d-e) SPBCA reporting biotinylated (*) MMP-1(E200A) binding to collagen in the presence of increasing concentrations of MMP-1Cat, over 1.5 h at variable temperatures (d), or at 25 °C with variable incubation times (e). (f) SPBCA reporting biotinylated (*) MMP-1(E200A) binding to various collagens in the presence of increasing concentrations of MMP-1Cat, over 1.5 h at 30 °C. All SPBCAs were performed in a microtiter-plate format, using immobilized type I collagen purified from guinea pig skin (unless otherwise indicated), and using colorimetric detection of biotinylated proteins by absorption at 450 nm wavelength [A450] (for further details see Materials and Methods). Presented results are mean ± standard deviation (SD) of triplicate readouts, with no visible error bars for SD values smaller than data points.

Overall, the SPBCA results suggest that even at the top of the collagen physiological temperature range (37 °C), close to its thermal denaturation temperature, the ES_unw_ complex is still under strain. This means that the thermal plasticity of the substrate alone does not warrant the most optimal fit to the enzyme (maximal potential binding affinity). In other words, MMP-1 distorts collagen strands in a distinct manner, inducing strain in the ES_unw_ complex. The strain energy stored in the ES_unw_ complex cannot be released by simply supplying heat, unless collagen chains become completely denatured (above 37 °C) whereupon the whole complex dissociates. Conversely, the heat appears to enable the conformational change in the context of the enzyme-substrate interaction (ES_w_ → ES_unw_), likely by boosting thermal fluctuations (potential energy) in the complex. Thermal energy is, therefore, a sort of gatekeeper for the *unwinder* or triple-helicase [9] activity of MMP-1. The endothermic nature of the process is not surprising, given there is no other source of energy (such as, for example, hydrolysis of ATP) available to MMP-1 to drive collagen unwinding.

The apparent strain in the ES_unw_ complex gets readily released upon cleavage with MMP-1Cat added *in trans*, which presumably relaxes the otherwise strained collagen chain(s) in that complex, enabling tighter enzyme-substrate binding. Importantly, this cleavage is unnatural, as it pertains to the liberated collagen strand(s) other than the one docked in the active site cleft of the *unwinder* enzyme, which under natural circumstances would have been cleaved first. Crucially, the release of the strain through natural cleavage is expected to have the opposite effect from the one demonstrated for the unnatural cleavage, because the active site docking provides one of the multiple anchor points that maintain the strain (besides those in the Hpx domain known as *exosites*) [7], and it gets disrupted through the natural cleavage. Consequently, the natural cleavage in the ES_unw_ complex weakens the interaction with the now partially cleaved collagen, enabling a quicker processing of the remaining strands, which no longer requires input from the Hpx domain [3]. The unnatural cleavage was instrumental in revealing that, although the intrinsic affinity of MMP-1 for the unwound state (*K*_*unw*_) is roughly twice as high as that for the wound state (*K*_*w*_ ≈ ½ *K*_*unw*_), it is still suboptimal considering the potentially attainable (unconstrained) affinity, exhibited by the clipped complex. Therefore, the strain in the ES_unw_ complex is probably an important aspect of the mechanism of MMP-1, ensuring that the binding, after the initial cleavage, is not overly tight, allowing quicker dissociation of the enzyme and a faster turnover rate.

While the strong temperature-dependence of the interaction of MMP-1 with collagen shown here and elsewhere [7,33] can prompt a notion that the process is governed purely by conformational selection, there is also a substantial body of evidence, shown likewise here and elsewhere [3,14,15] that supports the active involvement of the enzyme in the catalysis of collagen unwinding. Unambiguous discrimination between the two classic models of dynamic macromolecular recognition mechanisms involving conformational changes: 1) selected fit (conformational selection) vs. 2) induced fit, is a common problem in biology, sparking controversies in different fields [34–38] but this may be a semantic, rather than a real issue, as most biological systems are not clear-cut cases. It has been argued that any given real situation, where *K*_*conf*_ or *K*′_*conf*_ ≠ 0, will be a combination of both mechanisms [26]. Accordingly, we postulate that both scenarios also account for collagen binding and unwinding by MMP-1. First, the enzyme clearly selects a particular, hydrophobic residue-rich and imino acid-poor collagenase-susceptible region, characterised by a less tightly folded triple-helix [10,11,16,39–42], as certain level of thermal plasticity in the substrate is, apparently, required for the efficient unwinding [11,16,17]. However, as previously argued [3] and shown here, MMP-1 is not just a passive acceptor of one or more of spontaneously fluctuating (stochastically sampled via so-called thermal breathing) unwound states of collagen, whose role is to merely stabilize those states (prolong their lifetime) and in turn increase the chance for an opportunistic cleavage in one of the separated collagen strands. Conversely, MMP-1 appears to utilise the pre-existing plasticity of the particular site in collagen (its intrinsic local property) to induce specific collagen unwinding, generating distinct strain in the unwound complex. This unique *unwinder* activity is fuelled by heat and compensated by entropy, which ensures sufficient structural dynamics in the enzyme-substrate complex (structural crosstalk) to trigger the unwinding machinery [15], in which the well-documented inter-domain flexing plays the key part [7,14,15,20,43,44].

## Acknowledgements

S.W.M. thanks Emeritus Professor Hideaki Nagase of Oxford University for mentorship, and Drs. Michelle Waniewski and Brittany Stinson from K.B.’s laboratory for guidance concerning ITC data collection. We thank Dr. Robert Visse of University of Oxford for purification of collagen I, Prof. Richard W. Farndale of University of Cambridge for the provision of the collagen peptide, and Prof. Stephen M. Krane of Harvard Medical School for the provision of mouse skin collagen and its non-cleavable version. S.W.M. was supported by the Kennedy Trust for Rheumatology Research.

## Author Contributions

S.W.M. designed and performed experiments and calculations, analysed data, interpreted results and wrote the manuscript; K.B. oversaw the research, interpreted results and edited the manuscript.

## Supplementary material

### Experimental Procedures

#### MMP preparations

ProMMP-1(E200A), proMMP-1Cat and proMMP-3Cat were overexpressed from a pET3a vector in *E. coli* BL21 (DE3) strain (Invitrogen). Transformed cells were grown to OD600 of approximately 0.4, then induced with 0.5 mM isopropyl-β-D-thiogalactopyranoside (IPTG) (Biogene), and harvested after 4 h. Inclusion bodies were collected by lysing the cells in 0.05 M Tris (2-Amino-2-hydroxymethyl-propane-1,3-diol)-HCl (pH 8), 0.1 M NaCl, 0.26 mg/ml lysozyme (Sigma-Aldrich), 1 mM ethylenediaminotetraacetic acid (EDTA), 0.5% Triton-X100 (BDH/VWR), and dissolved in 20 mM Tris-HCl (pH 8.6), 8 M Urea, 50 μM ZnCl_2_, 20 mM dithiothreitol (DTT) (BDH/VWR). The solution was then passed over a Macroprep HighQ ion-exchange column (BioRad), equilibrated in 20 mM Tris-HCl (pH 8.6), 8 M Urea, 1 mM DTT, 50 μM ZnCl_2_ (BDH/VWR) and eluted with a linear salt gradient (0-0.5 M NaCl). The fractions were run on SDS-PAGE and the peaks were pooled, diluted with 50 mM Tris-HCl (pH 8.6), 6 M Urea, 1 mM DTT, 150 mM NaCl, 5 mM CaCl_2_, 100 μM ZnCl_2_, 0.02 % NaN_3_ to A_280_ < 0.3, supplemented with 20 mM cystamine (Sigma-Aldrich) and refolded by dialyses at 4 °C against 4 volumes of renaturation buffer (50 mM Tris-HCl (pH 8.6), 150 mM NaCl, 5 mM CaCl_2_, 100 μM ZnCl_2_, 5 mM β-mercaptoethanol, 1 mM 2-hydroxyethyl disulphide, 0.02 % NaN_3_ (BDH/VWR)) for 24 h, then 10 volumes of the same buffer for another 24 h, then against 10 volumes of the same buffer without β-mercaptoethanol for 24h, and finally against 4 volumes of 50 mM Tris-HCl (pH 8.6), 5 mM CaCl_2_, 50 μM ZnCl_2_, 0.02 % NaN_3_, for 24 h. Refolded proteins were purified using Green A affinity column (Amicon), equilibrated with 50 mM Tris-HCl (pH 7.5), 75 mM NaCl, 5 mM CaCl_2_, 0.02 % NaN_3_, and eluted with linear salt gradient (0-1 M NaCl). ProMMPs were activated with MMP-3Cat in 50:1 molar ratio and 1 mM p-aminophenyl mercuric acetate (APMA) (ICN Biochemicals) in TNC buffer (50 mM Tris-HCl (pH 7.5), 150 mM NaCl, 10 mM CaCl_2_, 0.02 % NaN_3_) for 60-120 min at 37 °C. The mature MMPs were finally purified on an S200 gel filtration column (GE Healthcare) in TNC buffer.

#### Isothermal Titration Calorimetry

MMP-1(E200A) and a triple-helical collagen peptide (Ac-(GPO)2-GPO-GPQ-GLA-GQR-GIV-GLO-GQR-GER-(GPO)3G-NH2; O, hydroxyproline), synthesized in the laboratory of Prof. R. Farndale (Cambridge, UK) as described previously [7], were dialyzed extensively against 50 mM HEPES (2-[4-(2-hydroxyethyl)piperazin-1-yl]ethanesulfonic acid) buffer (pH 7.5), containing 150 mM NaCl and 10 mM CaCl_2_. We chose HEPES to reduce control experiments needed to remove the influence of the buffer in case protein-protein binding was coupled to (de)protonation of ionizable groups, as HEPES has small ionization enthalpy and heat capacity [45]. The collagen peptide (6 μM) was titrated with the MMP-1(E200A) (68 μm) at different temperatures using a MicroCal VP-ITC microcalorimeter. The instrument was programmed to carry out 15 injections of 10–20 μl each over 16 s, spaced at 300-s intervals. The stirring speed was 200 rpm. Heats of binding were determined by integrating the signal from the calorimeter, and binding isotherms were generated by plotting the heats of binding against the ratio of enzyme to substrate. The data were corrected for heats of dilution of MMP-1(E200A) and the Origin 5.0 software from Microcal Inc. was used to calculate the enthalpy changes (∆H) and stoichiometry (N).

#### Protein biotinylation

MMP-1(E200A) or type I collagen stocks (see further) were buffer-exchanged using Sephadex G-25M PD-10 desalting columns (GE Healthcare) into 50 mM N-Cyclohexyl-2-aminoethanesulfonic acid (CHES) (pH 8.8), 200⍰mM NaCl, 10⍰mM CaCl_2_. Then, 10⍰mM EZ-Link Sulfo-NHS-LC-Biotin (Thermo Fisher Scientific) solution in distilled water was added at 1:2 protein:biotin molar ratio and incubated for 1⍰h at room temperature. Proteins were next passed over another PD-10 column equilibrated in TNC buffer to remove excess biotin.

#### Solid-phase binding and cleavage assay (SPBCA)

Costar High Binding 96-well plates (Corning, UK) were coated with 50⍰μl of 20⍰μg/ml type I collagen in TNC buffer, incubated overnight at room temperature. They were then washed with TNC buffer containing 0.05% Tween 20 (TNC/T) and blocked with 3% bovine serum albumin (BSA) (Sigma-Aldrich) in TNC/T. The mouse collagens (wild-type and non-cleavable variant) were a gift of Prof. Stephen M. Krane (Harvard) and the guinea pig collagen was extracted by Dr. Robert Visse (Oxford, UK) as described previously [31]. Biotinylated MMP-1(E200A) (0.5 μM) was added in TNC buffer together with MMP-1Cat (1-5 μM) and incubated at 4–40⍰°C for 1-24⍰h. The wells were then washed briefly 3 times in TNC/T at the temperature of incubation and developed using streptavidin-horseradish peroxidase conjugate (R&D, UK) and 3,3⍰,5,5⍰-tetramethylbenzidine 2-Component Microwell Peroxidase Substrate Kit (KPL, UK) for a fixed time. All assays were carried out in triplicate and paired analyses were always developed simultaneously.

#### Western blot

SDS-PAGE gels were transferred onto a polyvinylidene fluoride (PVDF) membrane (GE Healthcare) at constant voltage of 25 V for 90 min in transfer buffer (20 % (v/v) methanol, 12.6 mM Tris, 96 mM glycine and 0.1 % SDS). The membrane was then blocked using 5 % BSA in TNC/T for 1 h at room temperature. After blocking, the membrane was washed for 10 min in TNC/T and then incubated for 1 h at room temperature in a solution of streptavidin conjugated with alkaline phosphatase (Streptavidin-AP) (Promega). The Streptavidin-AP was diluted according to the manufacturers’ instructions in TNC/T supplemented with 1 % BSA. The membrane was finally washed 3 times for 10 min in TNC/T and incubated with the Western Blue Stabilised Substrate for AP (Promega) until clear bands appeared. The membranes were scanned using Image Scanner TM III (GE Healthcare).

**Fig S1.**
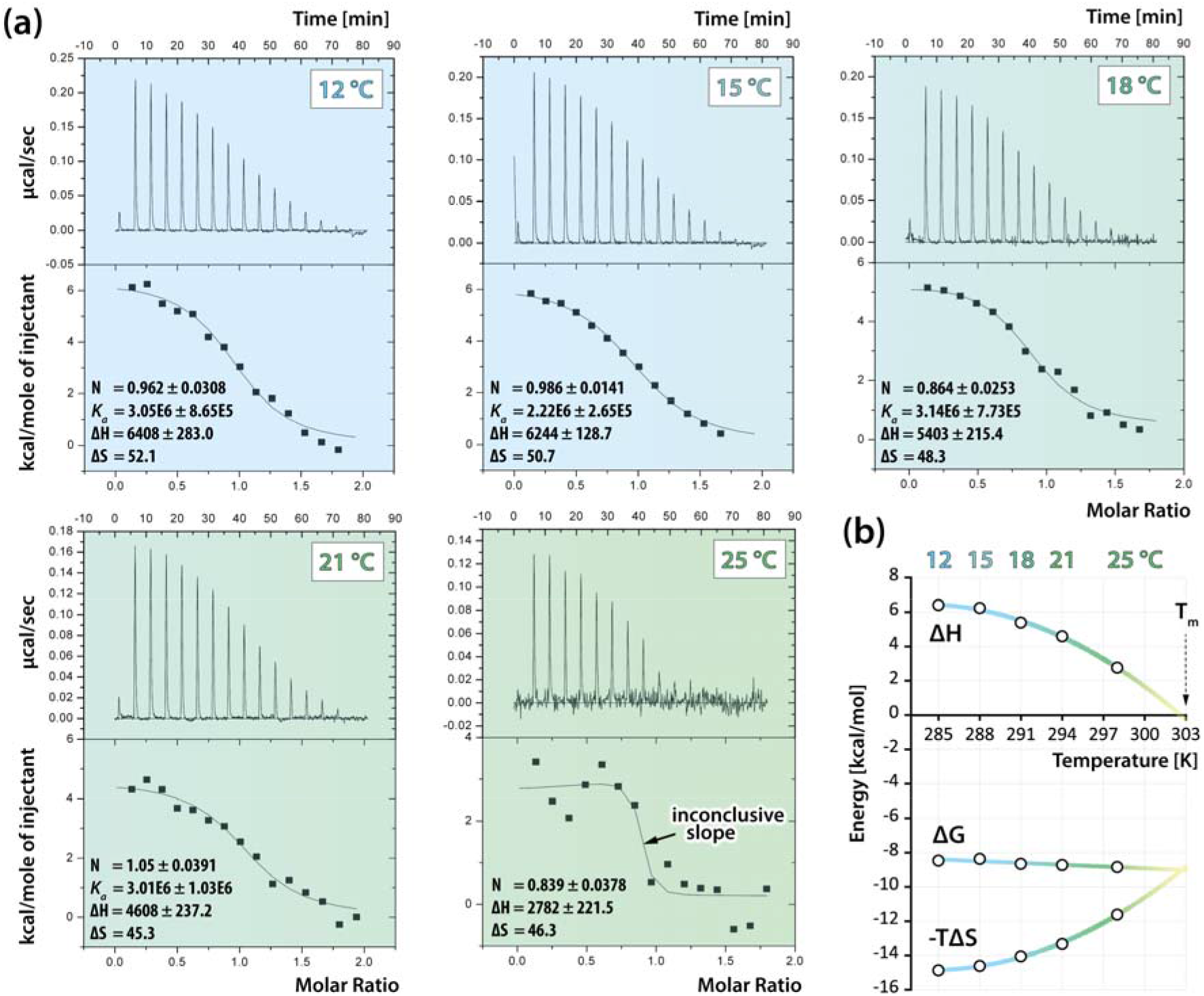
Isothermal calorimetric titrations of the triple-helical collagen peptide with MMP-1(E200A) at increasing temperatures. (a) Integrated heats from injections of equal volumes of the 68 μM stock of MMP-1(E200A) into 6 μM collagen peptide (Ac-(GPO)_2_-GPO-GPQ-G∼LA-GQR-GIV-GLO-GQR-GER-(GPO)_3_G-NH_2_; O, hydroxyproline; ∼, scissile bond), corrected for buffer-only injections (vehicle control), and fitted with isotherms using Origin 5.0 software (Microcal Inc.). Graph backgrounds are color-coded for temperature. Included are measured values of stoichiometry (N ≈ 1), enthalpy change (ΔH in kcal/mol) and equilibrium association constant (*K*_*a*_ in M^-1^, except for titration at 25 °C, due to inconclusive slope of the fitted isotherm). Included are also values of entropy change (ΔS in kcal/mol) calculated from the set of expressions for the Gibbs free energy change: ΔG = -RTln*K*_*a*_ and ΔG = ΔH - TΔS; R = 0.0019859 kcal/molK, gas constant; T, temperature in Kelvins [K]. (b) Thermodynamic signature of the MMP-1(E200A) interaction with the triple-helical peptide. ΔG and -TΔS terms were calculated as in (a) (average *K*_*a*_ from 12 – 21 °C range was used for the 25 °C data point). Fitted curves are color-coded for temperature and extrapolated until the peptide’s melting point (T_m_ ≈ 303 K or 30 °C).

**Fig S2.**
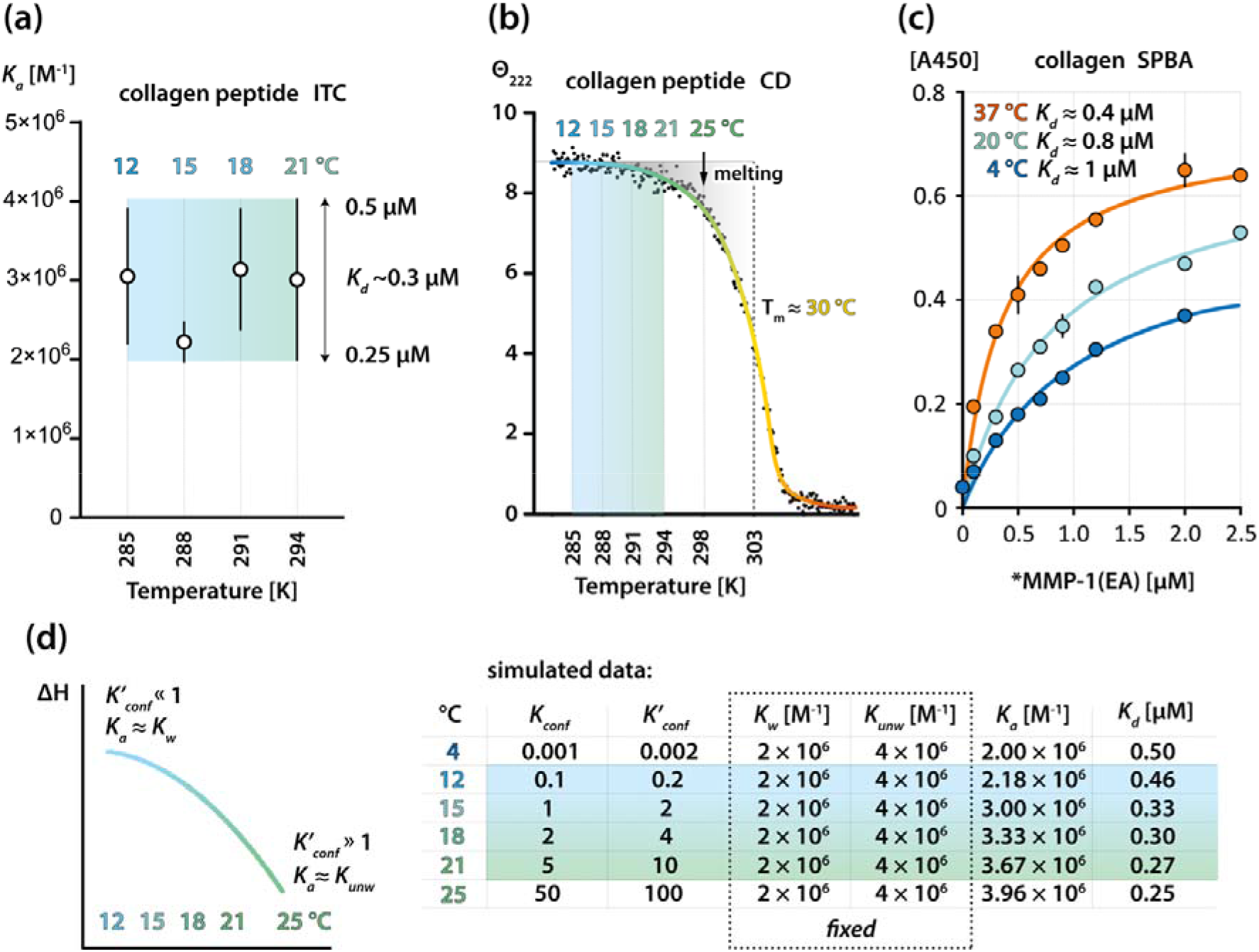
Temperature-dependence of the affinity of MMP-1(E200A) for the triple-helical collagen peptide. (a) A plot of equilibrium association constants (*K*_*a*_) of MMP-1(E200A) interactions with the collagen peptide measured at different temperatures with isothermal calorimetric titrations (ITC) (values ± error listed in Fig. S1a). The measurement uncertainty is demonstrated with the background shape color-coded for temperature and with the equilibrium dissociation constant (*K*_*d*_ = 1/*K*_*a*_) range. (b) Re-analysed data from [7]: melting curve of the collagen peptide (10 μM) obtained with circular dichroism (CD) ellipticity (Θ) at 222 nm. The fitted curve is color-coded for temperature and the reliable temperature range for direct *K*_*a*_ estimations by ITC is indicated with the likewise color-coded shading. (c) Re-analysed data from [7]: solid-phase binding assay (SPBA) reporting biotinylated (*) MMP-1(E200A) binding to immobilized collagen at increasing temperatures; [A450], absorption at 450 nm wavelength. Shown are the apparent *K*_*d*_ values estimated from the fitted curves for each considered temperature using Prism 8 (GraphPad). (d) Graph and data simulated to illustrate that the approx. 2-fold higher intrinsic association constant of MMP-1(E200A) for the unwound state (*K*_*unw*_), than for the wound state (*K*_*w*_) of collagen (as shown in (c)) will in the ITC experiment produce observed *K*_*a*_ (or *K*_*d*_) values falling within our measurement error. The observed *K*_*a*_ at each temperature is the population-weighted average of *K*_*unw*_ and *K*_*w*_: *K*_*a*_ = (*K*_*w*_ + *K*_*conf*_*K*_*unw*_)/(1 + *K*_*conf*_) or *K*_*a*_ = (*K*_*w*_ + *K*′_*conf*_*K*_*w*_)/(1 + *K*′_*conf*_(*K*_*w*_/*K*_*unw*_)), according to the conformational equilibrium constants (*K*_*conf*_ or *K*′_*conf*_) at each temperature: *K*_*conf*_ = [S_unw_]/[S_w_] = *K*′_*conf*_(*K*_*w*_/*K*_*unw*_); *K*′_*conf*_ = [S_unw_E]/[S_w_E] = *K*_*conf*_(*K*_*unw*_/*K*_*w*_). The table and the curve are color-coded as in (a) or (b).

